# Functional displacement of cortical neuromagnetic somatosensory responses: enhancing embodiment in the rubber hand illusion

**DOI:** 10.1101/2023.12.04.569949

**Authors:** Silvia L. Isabella, Marco D’Alonzo, Alessandro Mioli, Giorgio Arcara, Giovanni Pellegrino, Giovanni Di Pino

**Affiliations:** NeXT Lab, Campus Bio-Medico University of Rome, Rome, Italy; San Camillo IRCCS Research Hospital, Venice, Italy; Epilepsy program, Schulich School of Medicine and Dentistry, Western University, London, Ontario, Canada

## Abstract

The integration of an artificial limb as part of one’s body involves complex neuroplastic changes resulting from various sensory inputs to the brain. While sensory feedback is known to be crucial for embodiment, current evidence points merely to the attenuation of somatosensory processing, while the positive contributions of somatosensory areas to embodiment remain unknown. This study investigated the relationship between embodiment and adaptive neuroplasticity of early-latency somatosensory evoked fields (SEFs) in the primary somatosensory cortex (S1) following the Rubber Hand Illusion (RHI), known to induce short-term artificial limb embodiment. Nineteen healthy adults underwent neuromagnetic recordings during electrical stimulation of the little finger and thumb, before and after the RHI. We found a displacement of early SEF sources. In particular, we observed a correlation between the extent of rubber hand embodiment and specific changes to the m20 component (magnetic equivalent to the N20) in Brodmann Area 3b: a larger displacement and a greater reduction in m20 magnitude predicted the amount of embodiment, highlighting an important functional contribution of this first cortical input. Furthermore, we observed a posteriorly directed m35 displacement towards Area 1, known to be important for visual integration during touch perception (Rosenthal et al., 2023). Our finding that the larger displacement for the m35 did not correlate with the extent of embodiment implies a functional distinction between neuroplastic changes across these two components and areas in their contributions to successful artificial limb embodiment: (i) the earlier neuroplastic changes to m20 may shape the extent of artificial limb ownership, and (ii) the posteriorward shift of the m35 into Area 1 is suggestive of a mechanistic contribution to early visual-tactile integration that initially establishes the embodiment. Taken together, these findings suggest that multiple distinct changes occur during early-latency SEFs and their displacement in S1 last beyond the duration of the illusion and are important for the successful integration of an artificial limb within the body representation.

## Introduction

The sense that an artificial limb has become a part of one’s body involves neuroplastic changes arising from multiple and sometimes conflicting sensory inputs to the brain (Castro et al., 2023; Di Pino et al., 2009). However, many aspects of successfully embodying new body parts, or how this might fail (as occurs in up to 44%^1^ of amputees (Salminger et al., 2022)), remain largely unknown. Given that somatosensory feedback is crucial for embodiment (Cuberovic et al., 2019; Di Pino et al., 2020; Fritsch et al., 2021; Pinardi et al., 2020), this study sought to determine the adaptive processes within somatosensory responses that accompany embodiment of an artificial limb.

Since the primary somatosensory cortex (S1) is the first cortical region reached by tactile afferent inputs, it is thought to have a central role in the acceptance of a new body part into the body schema (Isayama et al., 2019; Shokur et al., 2013; Zeller et al., 2015). Current evidence suggests that S1 is attenuated in embodiment to facilitate prioritization of visual inputs (Castro et al., 2023). However, it remains unclear what are the *contributions* of S1 to the accommodation of an artificial limb into the body representation.

S1 is known to undergo significant neuroplasticity in response to use or environmental changes (i.e., bottom-up processes), balanced with changes in expectations (i.e., top-down processes) (Savolainen et al., 2011). Neuroplastic cortical remapping following loss of function, such as losing a body part, is well-documented (e.g., (Dykes & Metherate, 1988; Kaas et al., 1983; Makin et al., 2015; Pellegrino et al., 2012)). One way to investigate this type of remapping employs peripheral electrical somatosensory stimulation, that causes a relay of activity along the somatosensory pathway towards the cortex (e.g., (Rossini et al., 1994)). Signals at specific latencies are known to result from activity within different structures. The first cortical signal after peripheral stimulation of the wrist at about 20 ms (N20 or m20 of the somatosensory evoked potential or field, SEP or SEF) reflects direct and indirect effects of thalamic input to Brodmann area (BA) 3b within S1 (Allison et al., 1989). Area 3b has relatively small and well-defined receptive fields, responsible for detailed analysis of tactile stimuli, and precise localization of the stimulus on the body map. Following this, outputs from BA 3b and the thalamus arrive to BA 1 as early as 25 ms (P25), in which there are larger receptive fields that integrate sensory inputs from adjacent areas of the body, for complex perceptual judgments such as object recognition (Besle et al., 2014; Burton & Fabri, 1995; Macerollo et al., 2018; Martuzzi et al., 2014). Interestingly, it has recently been found that BA1 was more responsive to tactile stimulation when visual information was included (Rosenthal et al., 2023). While it is known that visual information is essential for artificial limb embodiment (Tsakiris & Haggard, 2005), the study by Rosenthal and colleagues points to the specific involvement of BA1 in visuo-tactile integration for successful embodiment.

The most-studied cortical responses to somatosensory stimulation occur at early (20-50ms) and mid-latencies (50-100 ms). Although the effect of embodiment on displacements of the early-latency components has not been studied, there are interesting findings for the mid-latency component which reflect the interaction between several brain regions that convey complex stimulus information and are generally considered to represent higher cognitive processes in comparison with shorter-latency responses (Desmedt et al., 1983; Hillyard & Kutas, 1983; Wu et al., 2012). Among these mid-latency responses, an important study found evidence for neuroplastic cortical remapping following the addition of a body part: displacement of the m60 source was observed during the illusion of having a third arm (Schaefer et al., 2009). However, effects of embodiment on displacements of early-latency S1 responses remain to be determined.

Early-latency cortical responses related to stimulus processing are linked to the inflow of sensory information, and they are subject to state dependant changes. For example, a displacement of the source of m20 activity as well as the source at about 50 ms (m40) (BA 3b and 1, respectively) was observed following disuse by anesthesia (Rossini et al., 1994). The displacement of the m40 was similarly observed due to long term use in violin-players (Elbert et al., 1995). This use/disuse displacement in early SEF sources is a neuroplastic change that may be an important mechanism underlying changes to the body representation. These early-latency responses (20-50 ms), reflecting initial stimulus processing, have been shown to contain most of the clinically relevant cortical somatosensory response components within S1 (Carter & Butt, 2001; Mauguière, 2003). For example, only components within these latencies are impacted by astereognosis (inability to identify objects by touch; (Mauguière et al., 1983)), and thus will be the focus of the current study. Displacements at the earliest latencies could point to a functional contribution of basic fundamental somatosensory processes (e.g., stimulus encoding) within S1 to successful embodiment.

To study the integration of an artificial limb within the body representation and its neural correlates, the Rubber Hand Illusion (RHI) is often employed in healthy adults and in amputees (D’Alonzo et al., 2015; Ehrsson et al., 2008; Schmalzl et al., 2014). In this paradigm, the participant experiences an illusion of owning the fake hand (i.e. embodiment) that begins within seconds. The illusion occurs when the participant observes an artificial rubber hand stroked with a paintbrush by an experimenter, who synchronously strokes the subject’s hidden real hand (Botvinick & Cohen, 1998; Ehrsson et al., 2004; Lloyd, 2007). This illusion is thought to arise from the complex integration of bottom-up multisensory information and top-down expectations about sensory information (Armel & Ramachandran, 2003).

The effects of the RHI on S1 and related somatosensory areas have been studied by measuring SEPs using electroencephalography (EEG), yielding mixed results. Several studies have found that the RHI enhanced a long-latency component at 140 ms (N140) (Kanayama et al., 2007; Press et al., 2008), likely originating in the secondary somatosensory cortex (Hari et al., 1983). One study demonstrated that the RHI reduced a mid-latency component around 50 ms (P45) (Zeller et al., 2015). Lastly, only one study demonstrated effects within the early-latency components: the RHI reduced activity within the 20-25 ms time window (N20-P25 component, occurring in BA 3b and 1, respectively (Sakamoto & Ifuku, 2021)). Although this study demonstrates a relationship between the RHI and the earliest SEPs, it remains unknown whether this attenuation of activity is accompanied by any adaptive processes to accommodate the artificial limb.

Early SEPs or SEFs provide accurate information on the location of sensory stimuli on the body, and a relative shift in SEF source location could be an adaptive process enabling successful embodiment. That is, a change in SEF source location following the RHI could underlie changes to the body schema, as might occur after loss or gain of a body part. Furthermore, it is known that this illusion relies upon top-down processes to prioritize visual somatosensory information (Taskiris & Haggard, 2005). Therefore, the question driving the current study is whether there is a relationship between embodiment and effects on somatosensory representation areas within S1. We expect to observe stronger changes in BA1, occuring *after* the earliest components at 20 ms within BA 3b, thus representing embodiment effects at the onset of integrative processes within sensory cortices, that coordinate a reduction in representation of neighbouring body parts.

Most previous studies investigating neuroplasticity associated with the embodiment of an artificial limb have relied upon EEG that has insufficient spatial resolution to observe shifts in SEP sources, or functional magnetic resonance imaging (fMRI) that has insufficient temporal resolution. Thus, the current study was conducted using magnetoencephalography (MEG), having millisecond-temporal and millimetre-spatial resolution (Hämäläinen et al., 1993; Hedrich et al., 2017) to determine the relationship between the RHI and early-latency SEF source locations within S1. Subjects underwent neuromagnetic recordings during electrical stimulation of the little finger and thumb immediately before and after the RHI, to quantify changes in SEF components and sources due to artificial limb embodiment. The extent of embodiment was measured using validated questionnaires (Botvinick & Cohen, 1998), and correlated with neuromagnetic findings.

## Materials and Methods

### Participants

Nineteen healthy adults (13 females, range 22-56 years) participated in this experiment. All volunteers signed a written informed consent before their participation in this study. The study was approved by the local Ethics Committees (Province of Venice and Campus Bio-Medico University of Rome) and the protocols are in accordance with the Declaration of Helsinki and future amendments. All 19 subjects complied with task instructions and completed the experiment. One subject (male) was excluded from analyses due to abnormalities discovered in the structural MR image. Data from the remaining 18 subjects were analyzed.

### Experimental Procedure

The experimental procedure lasted about 30-minutes, and was performed as follows. All subjects completed the Edinburgh Handedness Inventory (Oldfield, 1971) to assess their hand dominance. Subjects sat upright in a comfortable armchair inside a magnetically shielded room with their eyes closed. For SEF assessments, ring electrodes were placed on the little finger and thumb (Little and Thumb, respectively) of the left hand for each subject and session during neuromagnetic recording. Using a high voltage stimulator (DS7A, Digitimer Ltd., UK), 400 stimulus repetitions (200 per finger) were delivered with the following parameters: square pulse duration 0.2 ms, interstimulus interval ranging from 250 to 270 ms and amplitude 3-times the sensory threshold. Stimulation to the little finger and thumb was randomly interleaved, with the stimulator housed outside of the magnetically shielded room.

This was conducted before (Pre) and immediately after (Post) a five-minute synchronous RHI procedure (**Figure 1A**). For the RHI, subjects were instructed to place their left hand on a wooden platform, with the upper arm and shoulder covered by a towel. While the left hand was occluded from view by a vertical panel, a lifelike rubber hand was positioned next to the panel in a visible position, at a distance of 15 cm from the subject’s left hand. Both the real (left) and rubber hands were fitted with nitrile examination gloves (**Figure 1B**), to make the hands look similar. The experimenter instructed the participants to fixate on the artificial hand for the entire duration of the RHI procedure. During the 5-minute procedure, the experimenter stroked the participant’s left and the rubber hand with two identical paintbrushes at a pace of approximately 1Hz.

**Figure 1.**
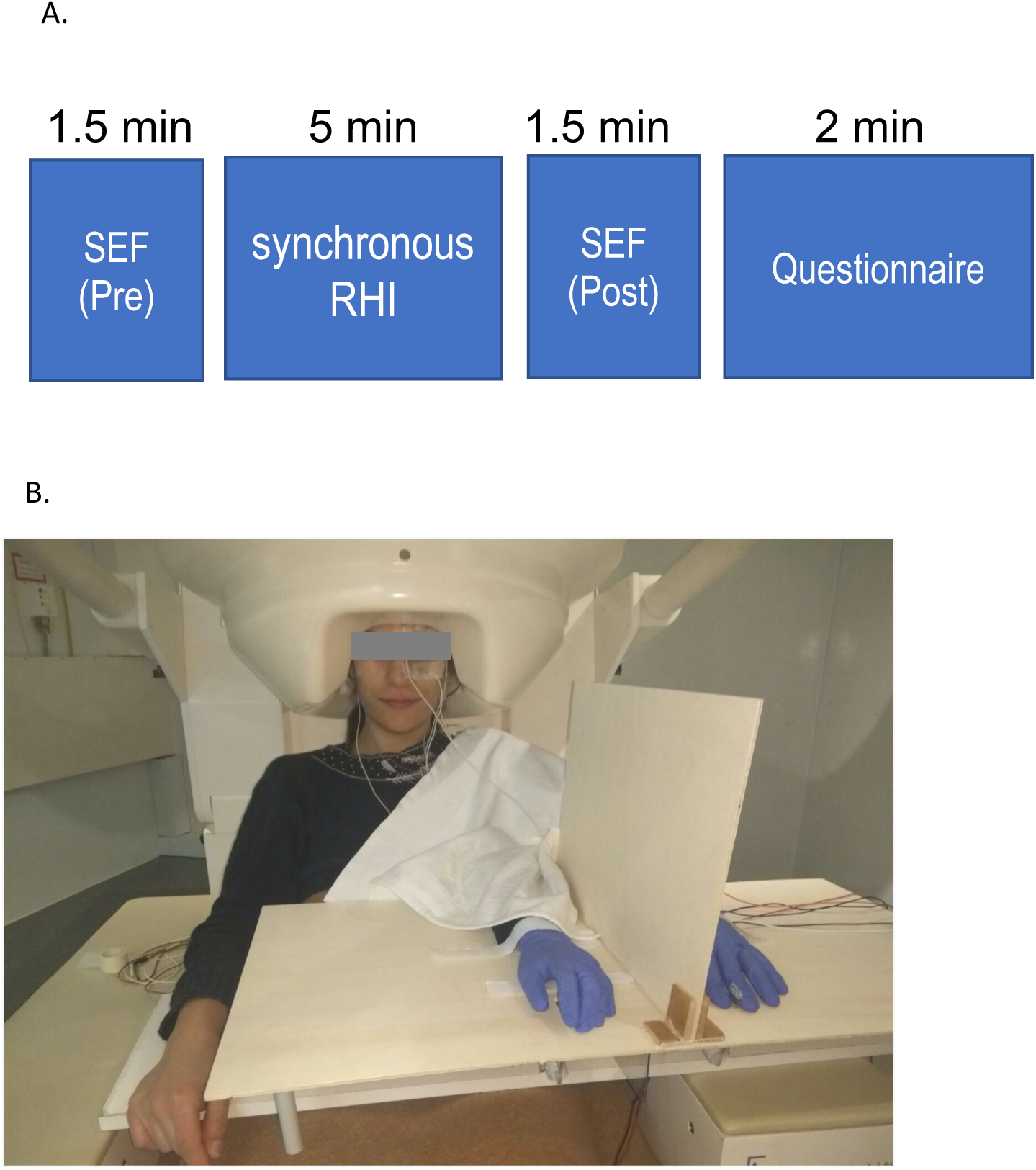
Experimental procedure and Setup. (**A**) Diagram of experimental procedure. Neuromagnetic evoked activity of thumb and little fingers electrical stimulation was collected before (Pre) and after (Post) the administration of synchronous RHI that lasted 5 minutes. The extent of embodiment was quantified through the RHI-index calculated from the answers to a questionnaire administered immediately following the procedure. (**B**) Rubber hand illusion setup, occurring between Pre and Post SEF procedures.

### Behavioural Measures

To quantify the extent of self-attribution to the rubber hand, participants were provided with a nine-item questionnaire ((Botvinick & Cohen, 1998); Supplementary Material) with which participants were asked to rate the extent to which the nine items did or did not apply, using a 7-point scale. For this scale, -3 meant “absolutely certain that it did not apply,” 0 meant “uncertain whether it applied or not,” and +3 meant “absolutely certain that it applied.” In order to control for participant suggestibility, three items in the questionnaire measure the illusion, whereas the other six items served as control for compliance, suggestibility, and “placebo effect”. From this, the RHI-index is calculated as the difference between the mean score of the illusion items compared with the mean score of the control items (Abdulkarim & Ehrsson, 2016; D’Alonzo et al., 2019), and serves as a quantification of the extent of embodiment experienced for each subject and condition.

### Magnetoencephalograph Recordings

Neuromagnetic activity was recorded using a whole-head 275-Channel CTF MEG system (VSM MedTech Systems Inc., Coquitlam, BC, Canada) in a magnetically shielded room. Data were collected at a rate of 1200 samples/s. Small coils placed at fiducial locations (nasion and preauricular points) were used with continuous head localization to monitor head position during recording. In order to localize MEG activity to each individual’s anatomy, T1-weighted structural MR images were collected for each subject using a 3T Ingenia CX Philips scanner (Philips Medical Systems, Best, The Netherlands). Head shapes and fiducial locations were digitized using a 3D Fastrack Digitizer (Polhemus, Colchester, Vermont, USA), which was used to co-register source images to the subject’s MRI using the Brainstorm Matlab toolbox (Tadel et al., 2011).

### Magnetoencephalograph Preprocessing and Source Analysis

Continuously recorded MEG data were segmented into 200 epochs of 1500 ms duration (500 ms pre-stimulus baseline), each for Little and Thumb, and for each session (Pre and Post). Epochs in which the peak-to-peak amplitude across MEG channels exceeded 3 pT during the time points of interest (-50 to 240 ms) were automatically labeled and rejected following visual inspection, resulting in a mean of 170 epochs (SD = 27.7) for each finger and session included for analysis. Due to excessive noise for one subject, data from 120 channels were excluded, and analysis was performed on the remaining 153 channels for that subject.

Data were highpass filtered off-line at 0.01 Hz, with a band-stop filter (50, 100, 150 Hz) to remove power line noise. Mean head position was calculated offline, and a multi-sphere head model (Lalancette et al., 2011) was used, implemented in the BrainWave Matlab toolbox (Jobst et al., 2018). In order to measure any differences in source location of SEFs, an event-related beamformer (Cheyne et al., 2006, 2007) with 2 mm spatial resolution was used to generate source activity images for averaged brain responses. This is a spatial filtering method that computes volumetric images of instantaneous source power corresponding to selected time points in the average (evoked) brain responses.

Data from Pre and Post sessions were combined for each finger to compute the data covariance used in estimating the beamformer spatial filter weights from the single trial data. In order to exclude effects from the stimulus artifact, the covariance window used was 10-240 ms from stimulus onset (Cheyne et al., 2007). Beamformer images were created every 1 ms over the period between 15 and 50 ms following the stimulus. Subsequently, peaks of activation across space and time were determined within this time window, and source direction was aligned across subjects, in native source space. This process was repeated for identified peaks of descending magnitude, until a discrete peak for each condition could no longer be identified. Given that, for example the m20 component may occur at slightly different latencies and source coordinates between subjects, a *component* is the label given to the peaks of activation across conditions for a between-subject average source and latency (e.g., *the m20 component*). For group averaging, spatial normalization was based on the MNI (T1) template brain, and subsequent scaling to Talairach coordinates were carried out using SPM12 (Wellcome Centre for Human Neuroimaging, London, United Kingdom). MNI coordinates of group-averaged source locations were plotted onto the ICBM152 template brain using Brainstorm.

### Statistical Analysis

For all acquired MEG and behavioural data, normality was tested using the Shapiro-Wilk test. If the data were not normally distributed, non-parametric statistical tests were used.

In order to confirm the induction of the illusion, we calculated the RHI-index as the difference between the mean scores of the illusion items and the mean scores of the control items. This index was used as the illusion outcome for the RHI condition.

To verify that the results of the RHI questionnaire was not due to participant suggestibility, the mean score of the three items employed to measure the illusion was compared with the mean score of the six items that served to control for compliance, suggestibility, and placebo effect using a paired Wilcoxon Signed Rank test.

For each subject and session, source locations at different latencies of Little and Thumb were used to calculate their Euclidian distance in native space.

In order to determine the within-subject effects of RHI on the Euclidian distance between SEF source locations we conducted a 2-way repeated measures ANOVA with two factors (session (2 levels: Pre vs Post) and component (3 levels: m20, m35, m45)). For each component, planned Pre vs Post comparisons were conducted using paired t-tests comparing values.

The source activity for each component evoked by the electrical stimulation was quantified. Since SEF peak magnitude values were not normally distributed, they were normalized using a cube-root transformation. In order to determine the within-subject effect of RHI on the SEF magnitude we ran a 3-way repeated measures ANOVA with three factors [session (2 levels: Pre vs Post), finger (2 levels: Little vs Thumb) and components (3 levels: m20, m35, m45)). For each component, planned Pre vs Post comparisons were conducted using paired t-tests comparing values.

In order to determine the presence of a relationship between embodiment measures and changes (Post-Pre) to evoked neuromagnetic activity, we conducted between-subject correlations: for each component (m20, m27, m35, m45) we correlated the change in Euclidian distance and the magnitude with RHI-index values using Spearman’s rank correlation. Although we did not hypothesize any specific relational differences between fingers, and therefore averaged magnitudes for Little and Thumb, post-hoc comparisons considered them separately. For the m27, only magnitudes for Thumb was used. All statistical tests were conducted and plots were prepared using R Statistical Software (Team, 2017). Corrections for multiple post-hoc comparisons were performed using Holm-adjusted values.

## Results

### Rubber hand illusion index

For the RHI (conducted between Pre and Post SEF procedures), mean RHI-index values were 4.19 ± 0.38 (Figure 2), which are comparable to values obtained in previous studies (e.g., (D’Alonzo et al., 2020)), confirming induction of the illusion. The mean value of the illusion items was significantly higher (2.4 ± 0.15) than the mean value of the control items (-1.75 ± 0.31; *p* < 0.001; Figure 2). Thus, the occurrence of the RHI cannot be attributed to the suggestibility of the participants.

**Figure 2.**
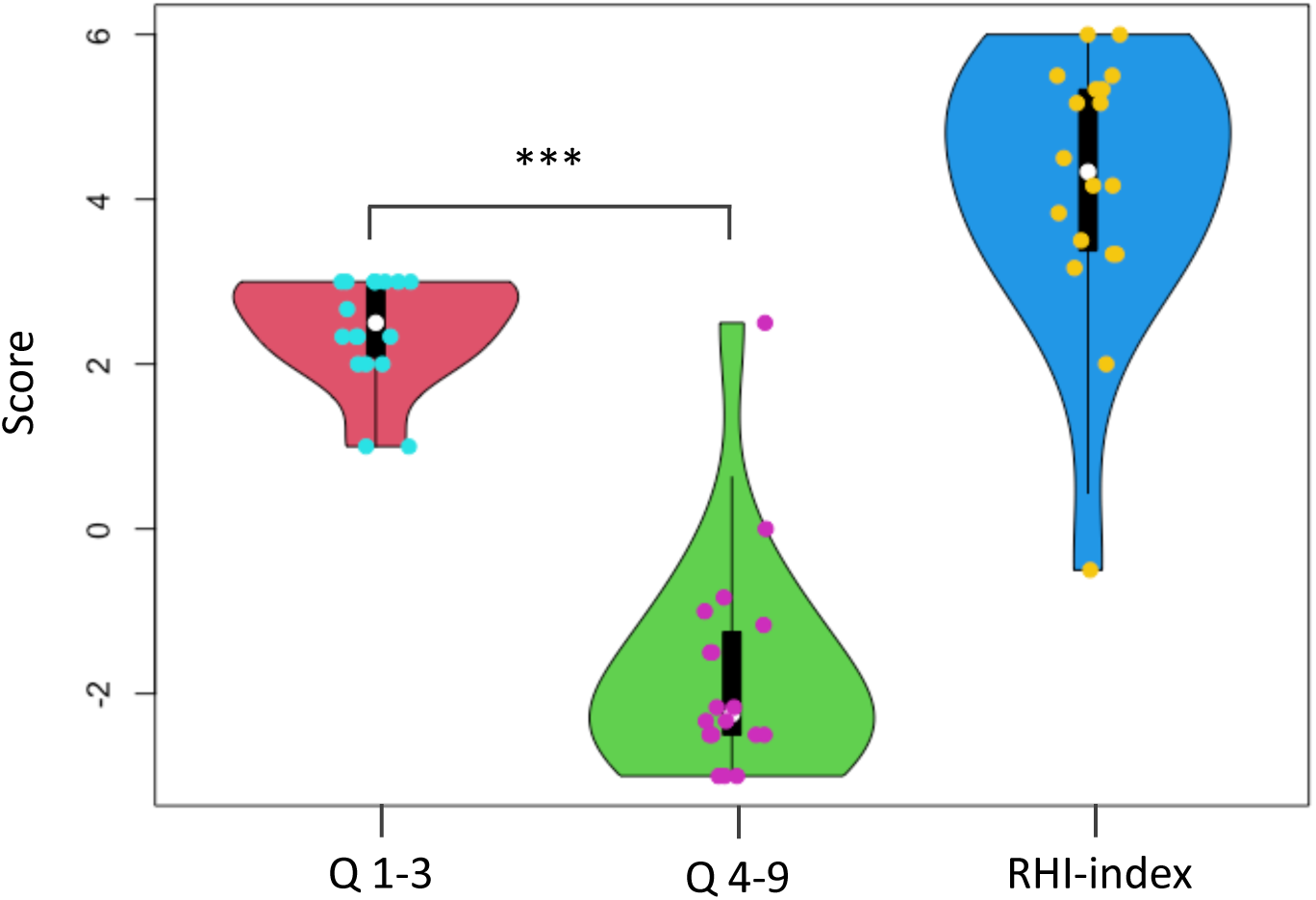
Mean values obtained for the RHI questionnaire. Questions 1-3 were the illusion-related questions, while questions 4-9 were the control questions. The significant difference between these sets of questions (*** = *p* < 0.001) demonstrate that the illusion was not merely due to participant suggestibility. The RHI-index quantifies the extent of the illusion, and values are comparable with those obtained in previous studies.

### Somatosensory evoked fields: within subject comparisons

Source-level analysis revealed peak averaged activity identified at 23 ms (m20), 28 ms (m27 – Thumb only), 34 ms (m35) and 44 ms (m45), the locations and latencies of which are summarized in **Table 1** and displayed in Figures 3 and **4**. Although the beamformer provided a clearly identifiable source of the m27 for Thumb (Pre and Post), there was no discrete m27 source for Little (Pre or Post) and therefore was not analyzed any further. This is likely due to the proximity of the m27 for Little to its m35 source (i.e., compare Figure 3B and C). For Little and Thumb, the m20 shifts were observed to be in opposite, anterior-posterior directions, whereas the m35 shifts were observed to mainly be in a similar, posterior direction (Figure 4).

**Figure 3.**
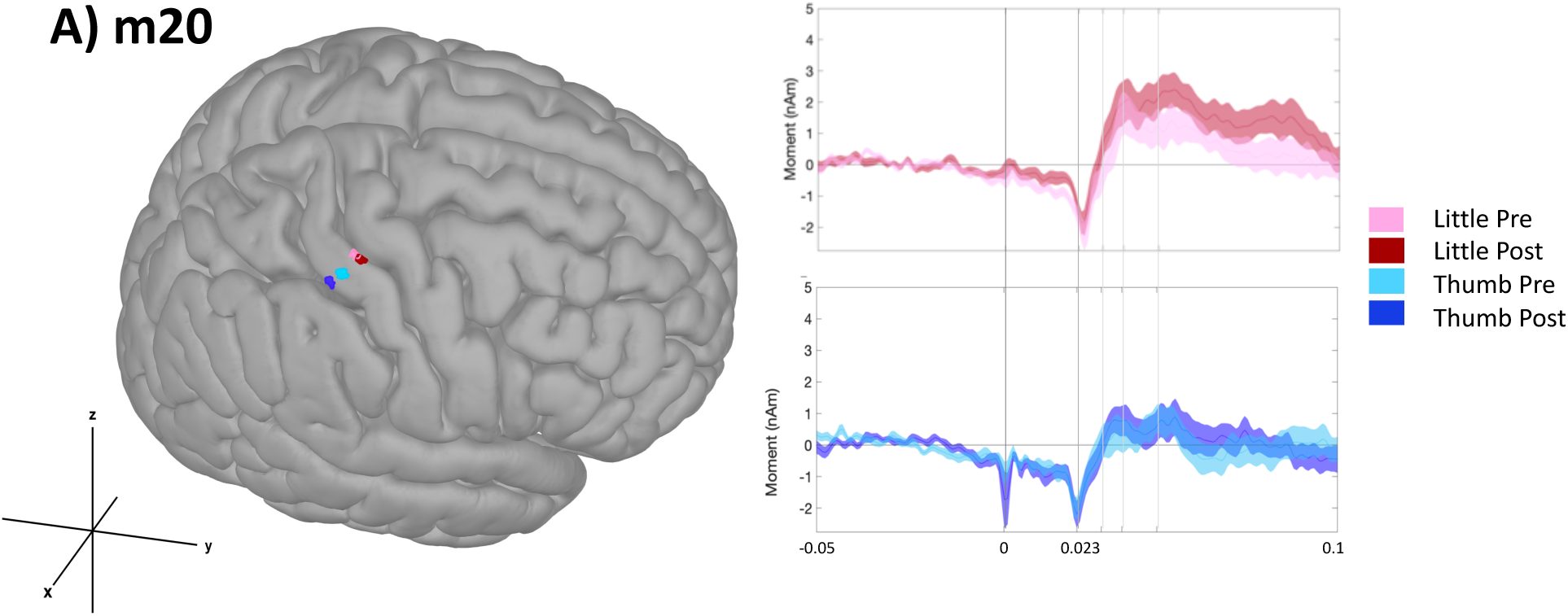
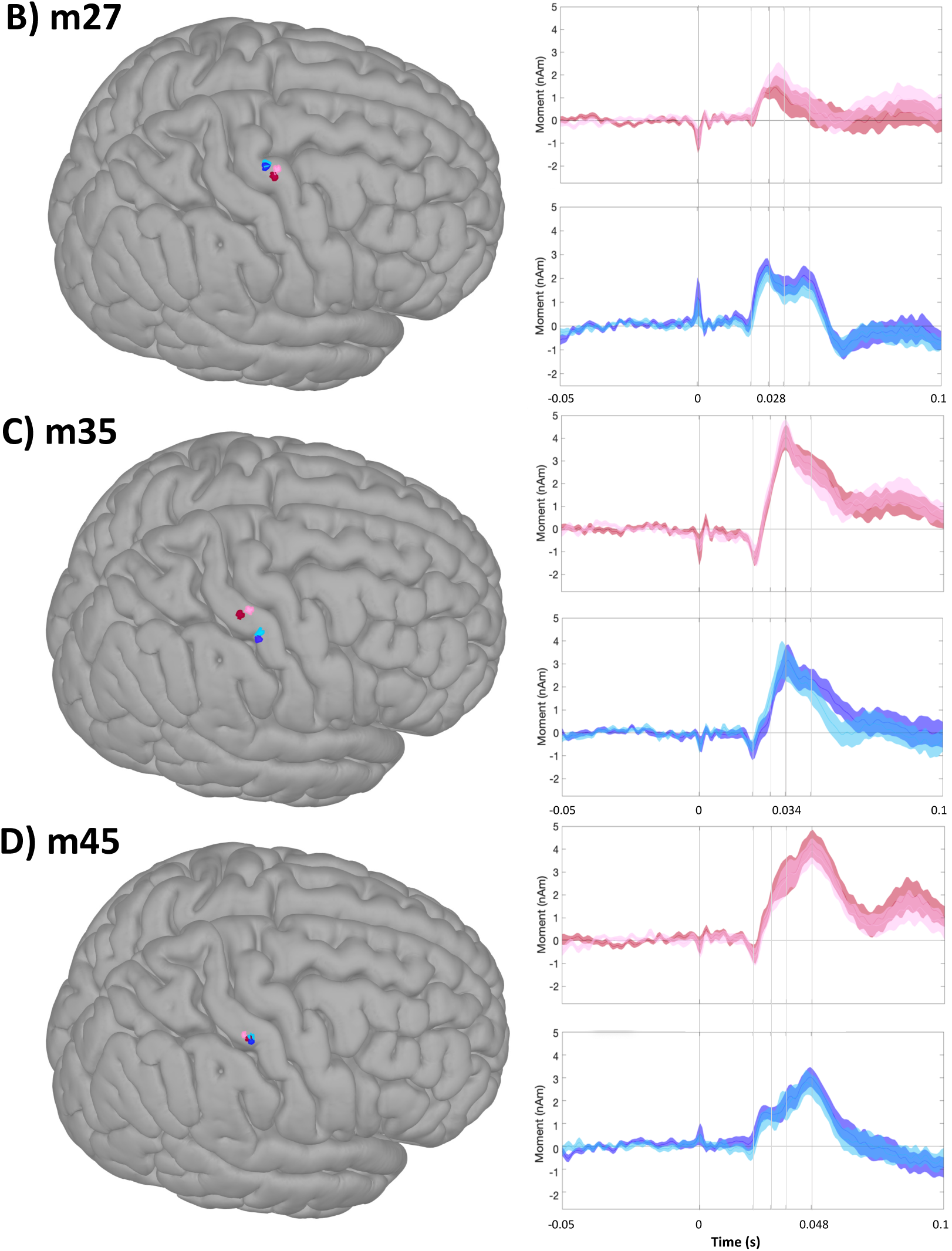
Mean SEF source location, plotted onto a template brain (left) and waveforms (right) for each finger, condition, and component. MNI coordinates are summarized in **Table 1**, and analyses are summarized in **Table 2**. **(A)** SEF location for the m20 component, with mean latency of 23 ms, localized to Brodmann area 3. **(B)** SEF location for the m27 component, with mean latency of 28 ms, localized to Brodmann area 6. Little sources were estimated and plotted for visualization purposes only, and not analyzed. **(C)** SEF location for the m35 component, with mean latency of 34 ms, localized to Brodmann areas 3 and 1. **(D)** SEF location for the m45 component, with mean latency of 44 ms, localized to Brodmann area 1.

**Figure 4.**
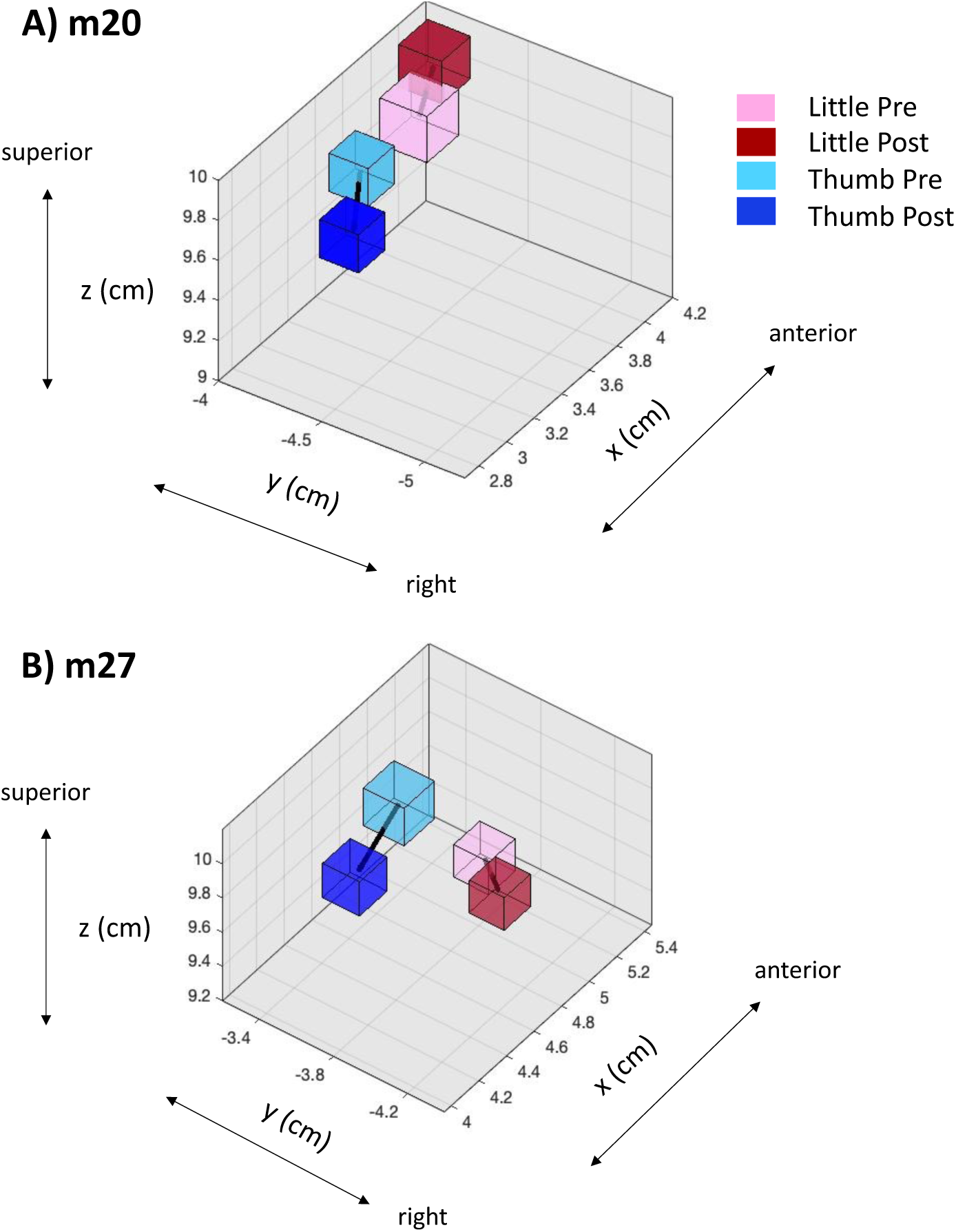
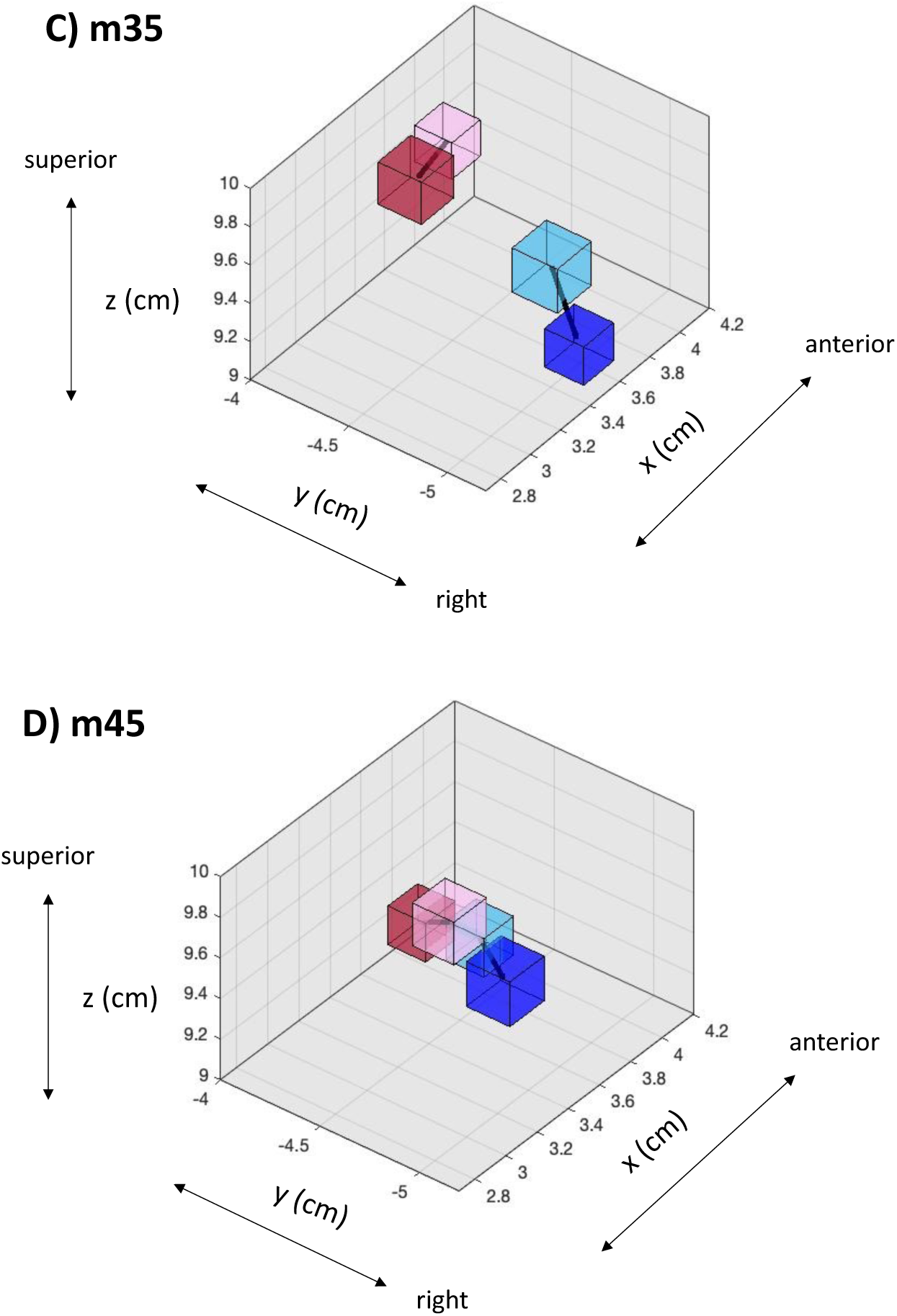
Mean SEF source location, plotted in native space for each finger, condition, and component. Shape volume corresponds to the standard error of the mean location. **(A)** SEF location for the m20 component, **(B)** m27 component, **(C)** m35 component and **(D)** m45 component. Note the change of axis values for **(B)** only, in the motor cortex.

**Table 1.** Mean somatosensory-evoked response component latencies and MNI coordinates for each finger and condition, where positive x-values denote right laterality, positive y-values denote anteriority, and positive z-values denote superiority. Analyses were conducted in native space. Note that a discrete source for the m27 component for Little stimulation could not be identified.

| Finger | Condition | Component | Latency (ms) | MNI coordinate |  |  |
| --- | --- | --- | --- | --- | --- | --- |
|  |  |  |  | X | y | z |
| Little | Pre | m20 | 23 | 46 | -21 | 66 |
|  | Post | m20 | 23 | 47 | -18 | 66 |
| Thumb | Pre | m20 | 23 | 50 | -25 | 62 |
|  | Post | m20 | 24 | 52 | -30 | 60 |
| Little | Pre | m27 | 30 | -- | -- | -- |
|  | Post | m27 | 30 | -- | -- | -- |
| Thumb | Pre | m27 | 28 | 40 | -12 | 72 |
|  | Post | m27 | 28 | 40 | -13 | 68 |
| Little | Pre | m35 | 34 | 50 | -22 | 62 |
|  | Post | m35 | 34 | 50 | -22 | 62 |
| Thumb | Pre | m35 | 35 | 58 | -20 | 58 |
|  | Post | m35 | 34 | 58 | -22 | 56 |
| Little | Pre | m45 | 44 | 54 | -24 | 60 |
|  | Post | m45 | 44 | 54 | -22 | 56 |
| Thumb | Pre | m45 | 44 | 54 | -24 | 60 |
|  | Post | m45 | 43 | 58 | -24 | 54 |

**Table 2.** Mean Euclidian distances between Little and Thumb SEF neuromagnetic sources, for each component. Note that m27 was excluded from statistical analyses, as described in the text.

| Latency | Euclidian Distance (cm) |  |
| --- | --- | --- |
|  | Pre (SD) | Post (SD) |
| m20 | 1.55 (0.59) | 1.82 (0.83) |
| m27 | - | - |
| m35 | 1.27 (0.52) | 1.63 (0.55) |
| m45 | 1.19 (0.60) | 1.23 (0.73) |

The analysis of the within-subject impact of RHI on SEF source Euclidian distances revealed the presence of significant main effects of session (Pre, Post: F(1,17) = 6.77, *p* = 0.019, *ηp*^2^= 0.28) and component (m20, m35, m45: F(1,17) = 7.612, *p* = 0.013, *ηp*^2^= 0.31), with no interactions (*p* = 0.62) (**Table 2 and** Figure 5). Planned comparisons revealed a trend towards an increase in Little-Thumb Euclidian distance for the m20 component (*p* = 0.11), reaching statistical significance for m35 (*p* = 0.021), but not for m45 (*p* = 0.42). Next, the repeated measures ANOVA conducted on peak magnitudes across components did not reveal a statistically significant main Pre-Post effect (F(1,17) = 0.011, *p* = 0.92).

**Figure 5.**
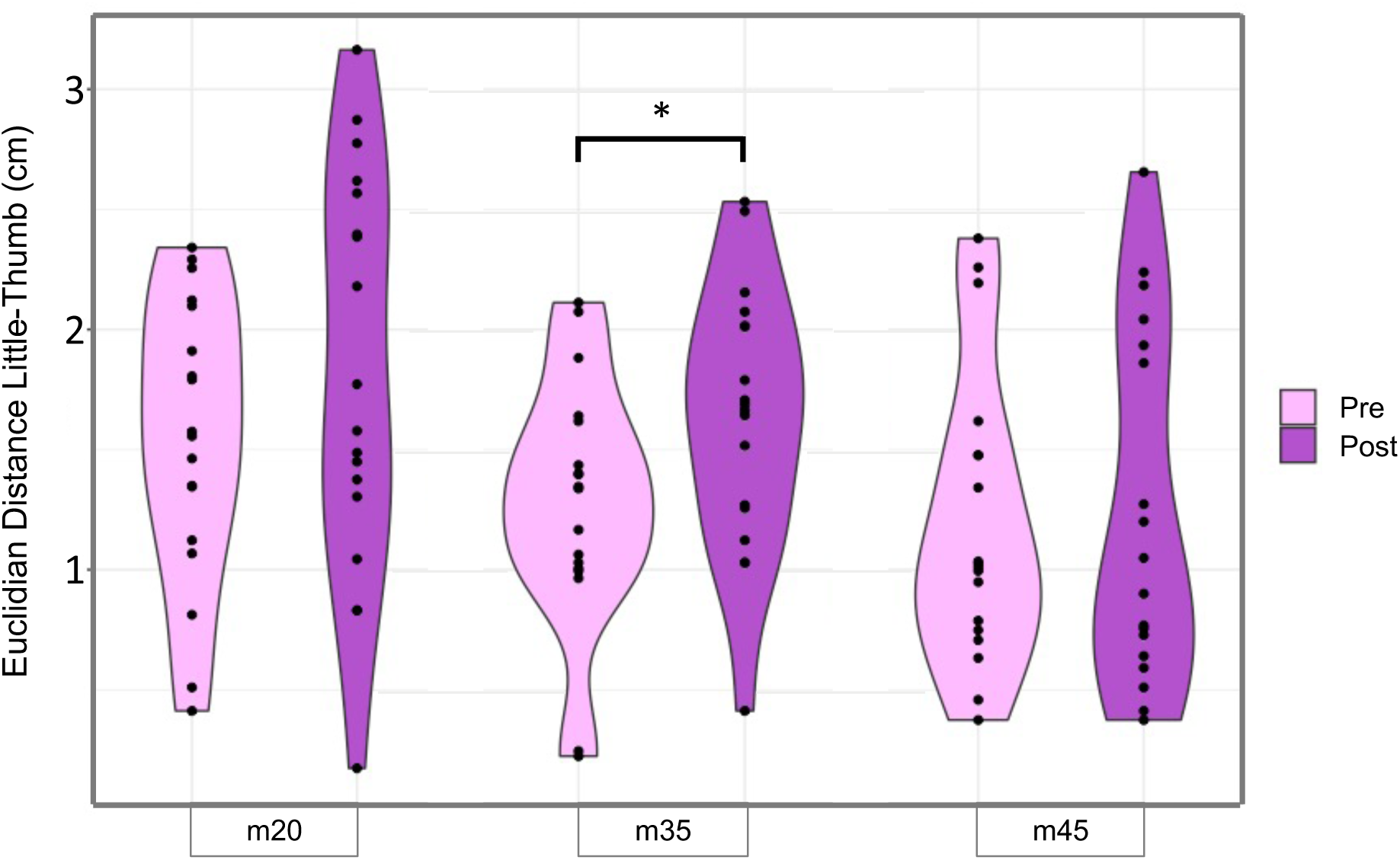
Mean Euclidian distances between Little and Thumb SEF neuromagnetic sources, for each component, with Pre on the left and Post on the right. Analysis of variance revealed a main effect of session (Pre vs Post: *p* = 0.019) and component (m20, m35, m45: *p* = 0.013). Planned comparisons revealed a significant increase in Euclidian distance for the m35 component (*p* = 0.021).

### Somatosensory evoked fields – rubber hand illusion-index: between subject correlations

In order to investigate between-subject relationships between the parameters chosen to quantify changes in S1 activity and the extent of embodiment achieved through the rubber hand illusion procedure, RHI-index was correlated with: (i) the change in Euclidian distance (with positive values indicating a greater distance after the RHI), and (ii) the average change in source magnitude (with negative values indicating greater reduction in signal strength after the RHI) for each component. For the m20, the Pre-Post change in Euclidian distance and magnitude correlated with the RHI-index (Euclidian distance: Spearman’s rho = 0.49 – moderate correlation, *p* = 0.02; peak magnitude: Spearman’s rho = -0.42 – moderate correlation, *p* = 0.04; Figure 6 and **Table 3**). No other planned correlations were significant. Post-hoc analyses revealed that the relationship between the RHI and the m20 magnitude was mainly driven by a reduction in activity for the Little finger (Spearman’s rho = -0.61 – moderate correlation, *p* = 0.008, < pHolm of 0.01), and not for the Thumb (Spearman’s rho = -0.01 – no correlation, *p* = 0.97). Given this finding, a similar post-hoc analysis for individual finger m35 magnitude revealed a relationship between the RHI and a decrease in m35 Thumb magnitude (Spearman’s rho = -0.61 – moderate correlation, *p* = 0.007, < pHolm of 0.008) was found, but not for Little (Spearman’s rho = 0.28, *p* = 0.26).

**Figure 6.**
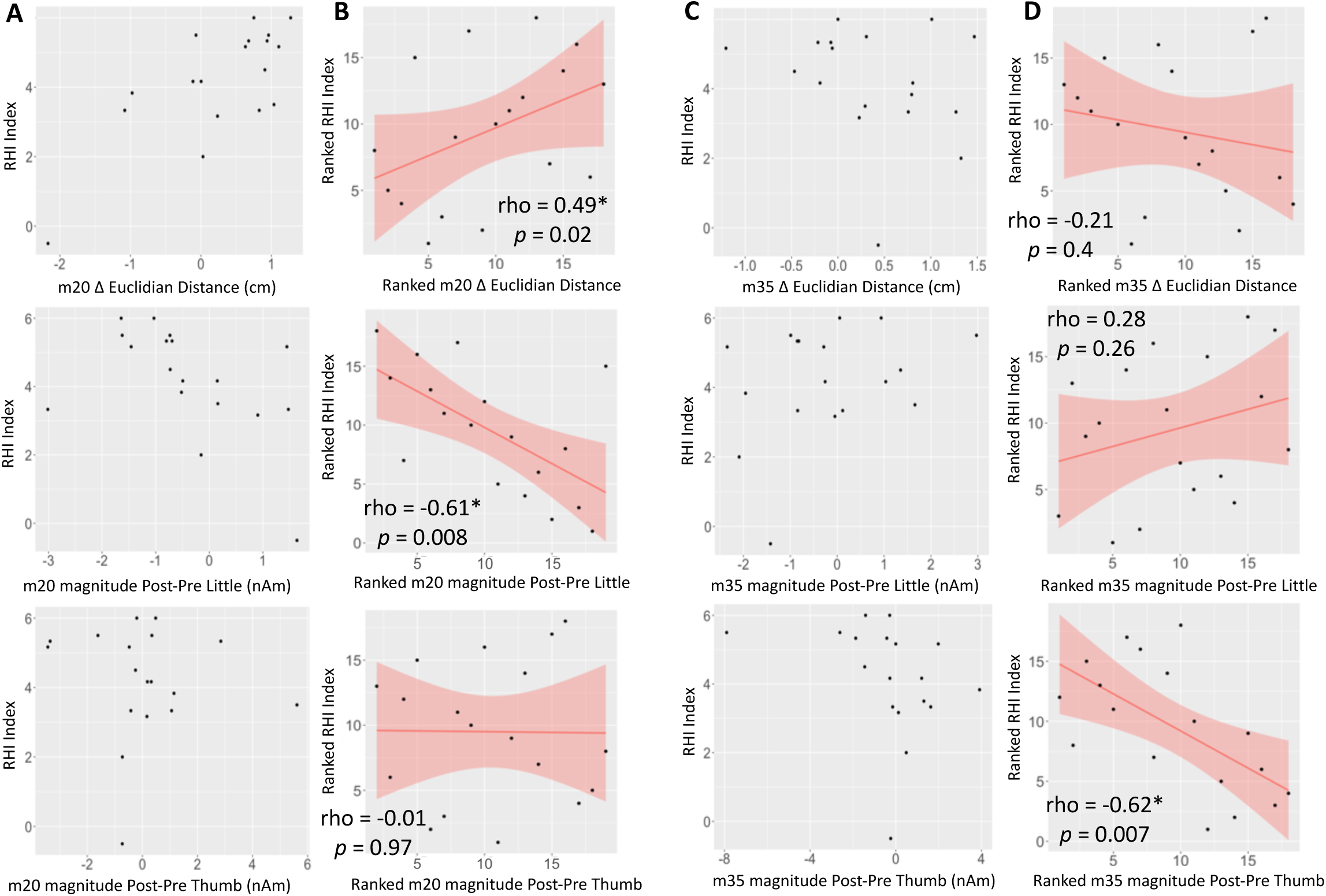
Comparisons between RHI-index and (A) m20 original values, (B) m20 ranked values, (C) m35 original values, and (D) m35 ranked values. Neurophysiological measures are presented as follows: (**Top**) Change in Euclidian Distance, Post - Pre; (**Middle**) magnitude for Little, Post - Pre; (**Bottom**) magnitude for Thumb, Post – Pre. Since the data are not normally distributed, the statistical tests were conducted using ranked data (**B** and **D**). Significant correlations were found between ranked RHI Index and ranked changes in m20 Euclidian distance (**B-Top**, Spearman’s rho = 0.49 – moderate correlation, *p* = 0.02), m20 Little magnitude (**B-Middle**, Spearman’s rho = - 0.61 – moderate correlation, *p* = 0.008), and m35 Thumb magnitude (**D-Bottom**, Spearman’s rho = -0.62 – moderate correlation, *p* = 0.007), where positive values denote *increases* after RHI. Significant correlations are marked with an asterisk.

**Table 3.**
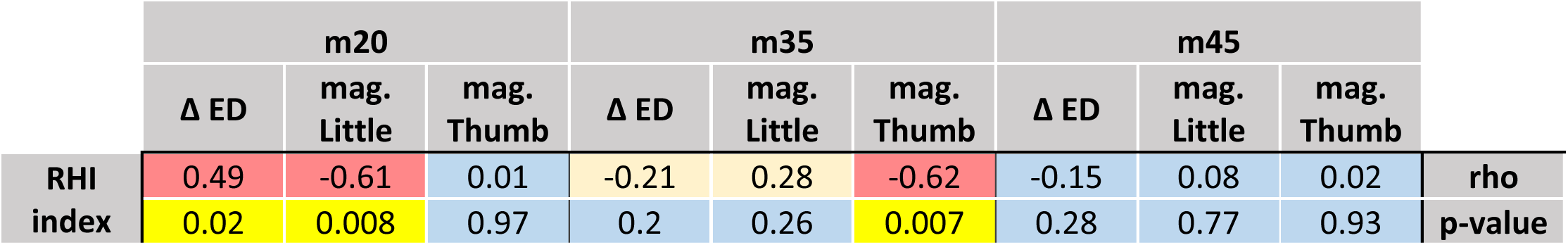
Matrix of Spearman’s rank correlations, comparing the RHI Index with the Pre-Post change in Euclidian Distance (Δ ED), and the average change in peak magnitude (mag.), as in Figure 6. Rho indicates the strength of correlation between ranked data, and the p-value indicates the significance of the correlation. Cells are colour-coded for strength of values (red for moderate correlation, light yellow for low, the p-value cell is yellow for significant correlation).

## Discussion

Most research to date on S1 activity during the RHI points to a reduction of SEPs, thought to reflect a prioritization of visual over somatosensory information. The current study investigated the early neuroplastic processes underlying how new body parts are accommodated by the sensory cortex. We observed a change in Euclidian distance between Little and Thumb sources across BA 3b and 1, with a statistically significant within-subject displacement occurring for the m35 component. These results support our hypothesis for a neuroplastic shift at latencies beyond the m20, reflecting integration processes, especially visuo-tactile integration known to be important for the RHI and occurring in BA 1 (Rosenthal et al., 2023; Tsakiris & Haggard, 2005). Furthermore, different neuroplastic shifts predicted the extent of rubber hand embodiment: between-subject MEG-illusion correlations of moderate strength were observed between the RHI-index (quantifying the extent of the illusion) and the m20 component within BA 3b across two parameters: (i) an increase in Euclidian distance, and (ii) the reduction in the magnitude of Little activity. An additional correlation was found between the RHI-index and the magnitude of Thumb activity for the m35 component. These differences across Brodmann areas, latencies, and fingers reveal a highly complex relationship between S1 and embodiment.

### The rubber hand illusion causes a shift in somatosensory evoked field source for early components

The effect of the RHI on source localization of early SEF components was of critical interest in the current study. The timing and locations of each SEF component are consistent with previous literature (Buchner et al., 1995; Rossini et al., 1994), including from invasive human recordings (Wood et al., 1988). That Thumb sources are represented relatively more posteriorly in the postcentral sulcus than Little is likely due to more distal positioning of the stimulating ring along the Thumb, as it has been shown that distal phalanxes are more posteriorly represented than proximal phalanxes (Roux et al., 2018). The Euclidian distances between Little and Thumb sources (averaged across subjects) ranged between 1.19 and 1.84 cm (**Table 3**), consistent with previous studies (Hari et al., 1993; Huber et al., 2020; Nakamura et al., 1998; Rossini et al., 1994; Wood et al., 1988).

We expected a change in Euclidian distance for the m35 component but not for the m20 component. This is because we were anticipating neuroplastic changes at the level of sensory integration that would furthermore correlate with the extent of embodiment. Although we expected this distinction between m20 and m35 components, it is very intriguing that the illusion measure correlated between-subjects only to the m20 displacement, and not the m35 displacement. It is notable that the magnitude of within-subject m20 displacement was nearly as large as the displacement for m35 (0.28 vs 0.36 cm, respectively), but with higher variance (mean SE 0.22 vs 0.16 cm, respectively). Thus, this variance that prevented within-subject statistical significance, that also *caused* the between-subject correlation, was related to differences in the extent of embodiment. This is an important consideration for future studies such as those using RHI that have high inter-individual differences.

Furthermore, the current results likely arise from a functional dissociation between the m20 and m35 components. That is, the SEF source displacement was larger (and more consistent) for the m35 than for the m20, but was not the displacement that correlated with the extent of embodiment. This additionally means that the stronger SEF source displacement for the m35 must have a different function from the relatively weaker SEF source displacement for the m20. As far as we are aware, these results provide the first evidence for multiple functional roles for SEF source displacements within S1.

These shifts are thought to exploit existing peripheral-S1 anatomical connections that remain functionally inactive under usual circumstances (Rossini et al., 1994; Sanes et al., 1988), similar to the early neuroplastic changes that are known to follow amputation (Di Pino et al., 2009). It is unknown whether there are any differences in the underlying mechanisms across m20 and m35 SEF displacements. The source of the m20 in BA 3b, the m27 in motor areas (BA 4 and 6), m35 in BA 3b and 1, and m45 in BA 1 is in line with known generators of these signals (Allison et al., 1989; Balzamo et al., 2004). With respect to the current findings that the m35 was localized either to BA 3b (Little) or BA 1 (Thumb), it is known that for this component, there is higher inter-individual variability and possibly multiple contributing sources (Huttunen et al., 2006; Kawamura et al., 1996; Peterson et al., 1995), likely reflecting a transition point with respect to inputs to BA 3b and 1. BA1 is known to receive direct thalamocortical inputs as well as inputs from BA 3b (Jones et al., 1978). Thus, the m35 component likely reflects a gradual transition point towards top-down processing.

The RHI is dependent upon the integration of visual, tactile, and proprioceptive information, which highlights the importance of top-down processes in shaping body ownership and embodiment. The observation that the m35 displacement is posteriorward towards BA 1 suggests the importance of visual-tactile integration in the embodiment of the fake hand (i.e., (Rosenthal et al., 2023)), which however is not directly related to inter-individual differences in the extent of embodiment. This is a rather intriguing finding, suggesting that this integration is not related to the extent of embodiment, but to some other component of the illusion, for example its initiation or its stability over time.

We did not expect to see shifts or correlations for the very first component (m20), as seen by Rossini and colleagues with decreased use (Rossini et al., 1994). Our finding that the m20 displacement correlated between-subjects with the extent of embodiment would suggest, beyond the expected effects of top-down integrative processes, that relatively more bottom-up processes related to low-level physical representation are central to embodiment in RHI. Although top-down processes are necessary for the RHI to take hold (Tsakiris & Haggard, 2005), these results suggest that processes preceding somatosensory integration may have an even more important role. The m20 component in area 3b is thought to reflect cortical relay encoding tactile features limited to individual digits, prior to integration across digits and across sensory modalities (such as with vision).

### Expansion versus contraction of cortical somatosensory finger representation

Given the theory that S1 must be suppressed to prioritize visual information in order for the RHI to occur (e.g., (Castro et al., 2023)), one might expect the Euclidian distances between digits to contract during the RHI. This is based on findings that somatosensory cortical plasticity related to disuse is typically associated with a contraction of unused digit representations (Elbert et al., 1998; Rossini et al., 1994) and increased use is conversely associated with an expansion of an overused digit’s representation (Elbert et al., 1995; Godde et al., 2003). However, it is possible that the RHI results in a contraction of somatosensory representation of the stimulated finger only, with a corresponding expansion of the ‘spared’ fingers (Feldman & Brecht, 2005). This is supported by the findings that for synchronously co-stimulated fingers, their cortical representations contracted, whereas for asynchronously co-stimulated fingers, their cortical representations were centred further apart (Braun et al., 2000; Pilz et al., 2004; Vidyasagar et al., 2014; Ziemus et al., 2000), as observed in the current study. Furthermore, while the RHI increases arousal (D’Alonzo et al., 2020; Di Pino et al., 2022), arousal in turn increases cortical inhibition, resulting in less overlap in cortical receptive fields, along with reduced strength of the thalamocortical connection (Castro-Alamancos, 2004). In this way, a stronger embodiment during the RHI may be related to a simultaneous increase in Euclidian distance between fingers not involved in the RHI, with a reduction in the strength of thalamocortical input, as observed in this study.

### Rubber hand illusion strength is associated with attenuated somatosensory evoked fields

Although we did not observe significant within-subject reduction in SEF magnitude for any of the measured components, we did observe unexpected differences in between-subject correlations of RHI index and individual digit magnitudes across m20 and m35. Both correlations were in the same direction, demonstrating that subjects who experienced a greater sense of embodiment of the rubber hand had a subsequent greater reduction in SEF magnitude, for the Little at m20 and for the Thumb at m35. We did not expect to find any differences across digits. A possible explanation for this finding comes from a recent study that observed within area 3b of S1 a homogeneous 3D structural architecture between digits 2-5, excluding the thumb (Doehler et al., 2023). Greater independence of the thumb has also been observed within area 3b of macaque monkeys (Lazar et al., 2023). It is unlikely that this independence holds for area 1, where integration across fingers is known to occur. Thus, it is possible that our findings of a relationship between embodiment and reduction in magnitude for the Little at the m20 within area 3b reflects this greater predisposition for inter-finger plasticity, excluding the thumb. Meanwhile, since integration is fundamental to area 1, it is unlikely that the thumb retains its independence, and thus thumb-related plastic effects may be more evident in this area during the m35.

These findings confirm and extend the main findings of previous studies, that measured the effect of the RHI on early SEP components and found a reduction in magnitude at similar latencies as the m45 measured in the current study (Zeller et al., 2015), and also when combining earlier latencies that span the 20-25 ms time range (Sakamoto & Ifuku, 2021), thought to originate across areas 3b and 1. Differences between the current results and those of the two aforementioned studies could be due to differences in recording methods, or differences in timing: the aforementioned studies recorded SEPs during the RHI, whereas in the current study the SEFs were recorded before and after the RHI. This is an important distinction, as it is possible that the reduction in SEF magnitude, especially for the m45 component observed by Zeller and colleagues, may be a transient effect, whereas the observations of this study might represent more stable effects of embodiment, lasting beyond the illusion itself.

### A note on the motor cortex: Inversion across the central sulcus

It was not surprising to observe a strong m27 component in precentral motor areas (BA 4 and 6), as this is a known SEF component. However, the discrepancy in source localization results between Little and Thumb responses was not expected. Given the estimated placement of the Little sources on the crown of the precentral gyrus, it is possible that the anatomical layout of the hand representation area results in the Little source being more radially-oriented for this specific component than the Thumb source, and therefore largely insensible by the MEG gradiometers. That said, it is intriguing that the estimated relative layout is in the opposite orientation with respect to the S1 sources. That is, the Little source is estimated to be *more lateral* than the Thumb source in the motor areas, but more medial than the Thumb source in S1. These findings are in line with traditional anatomical layouts of the sensory homunculus, as well as with recent updates to the motor homunculus, including a fMRI study demonstrating concentric medial-lateral representations for the fingers and thumb in the motor cortex, with the Little finger at the centre (Huber et al., 2020). Thus, it is not anatomically incorrect for the Thumb source to be medial to the Little source within the motor cortex, as it has representations both medial and lateral to the Little source. With respect to the double representation of the Thumb in the recent motor homunculus, what is perhaps most interesting about the current result is that the medial Thumb representation was stronger than the lateral representation, suggesting a functional difference between these motor cortical Thumb representation areas.

### Limitations

There are limitations to the current study. We did not measure SEFs immediately before and after RHI control sessions, as the induction of the observed effects to the SEF in the Post condition would invalidate any subsequent SEF procedures. Since it is known that repeated stimulation increases the magnitude of SEFs only at long latencies and does not affect the early components analyzed here (Dowman & Rosenfeld, 1985), we do not believe that our results are solely due to the repeated paintbrush stimulation during the synchronous RHI, especially since several findings in this study correlated with the RHI-index measure.

### Summary and Conclusions

The present study demonstrated that the RHI induced changes in the source localization of early SEF components in S1, distinguishing between components and suggested multiple functions of SEF source displacements. The extent of rubber hand embodiment was associated with a between-subject increased distance between Little and Thumb cortical somatosensory representation areas for the m20, and peak magnitude of the m20 and the m35 components for the Little and Thumb, respectively.

These findings contribute to our understanding of the neuroplastic changes underlying successful artificial hand embodiment and provide insights into its potential mechanisms, separate from the neuroplasticity induced by loss of function. Adaptations in S1 accompanying embodiment likely represent different and complex mechanisms compared with use and disuse.

Understanding the neuroplastic changes involved in artificial limb embodiment is crucial for the development of effective strategies to enhance the functionality and acceptance of prosthetic devices, and reduce the occurrence of phantom limb pain. Future research could further investigate the temporal dynamics of somatosensory processing during artificial limb embodiment and explore the relationship between neuroplastic changes and long-term prosthetic use and phantom limb pain, in amputees.

## Footnotes

1 The reasons for abandoning a prosthesis are many and varied. Lack of embodiment of the prosthesis is an important but not the sole factor for its abandonment. See (Murray, 2008) for a detailed discussion.

## Notes

### Competing Interest Statement

The authors have declared no competing interest.

