## Supplementary Table for "Functional displacement of cortical neuromagnetic somatosensory responses: enhancing embodiment in the rubber hand illusion"

**Supplementary Materials**

**Table 1**. List of items of the questionnaire. The first three (illusion) items refer to the extent of sensory transfer into the rubber hand and its self-attribution during the trial. The subsequent six (control) items serve as controls for compliance, suggestibility, and “placebo effect”. In addition, the vividness and prevalence statements are reported.

| **Questionnaire** | **Item** | **Rating** |
| --- | --- | --- |
| Item 1 | It seemed as if I were feeling the tactile stimulation at the location where I saw the visible hand touched | -3 – +3 |
| Item 2 | It seemed as though the stimulation I felt was caused by the touch on the visible hand |  |
| Item 3 | I felt as if the visible hand was mine |  |
| Item 4 | I felt as if the position of my real hand was drifting towards the visible hand |  |
| Item 5 | It seemed as if I had more than two hand or arm |  |
| Item 6 | It seemed as if the tactile stimulation I was feeling came from somewhere between my own hand and the visible one |  |
| Item 7 | I felt as if my real hand were turning ‘rubbery’ |  |
| Item 8 | It appeared as if the position of the visible hand was drifting towards my real hand |  |
| Item 9 | The visible hand began to resemble my own hand, in terms of shape, skin tone, freckles or some other visual features |  |
| Vividness | How realistic and life-like was the illusion that the visible hand was yours when it was experienced? | 1 – 9 |
| Prevalence | How long with respect to the length of section was the perception of this illusion? | 0 – 100% |
